# Anti-target chemical-space coverage of Indian medicinal plant phytochemicals

**DOI:** 10.64898/2026.10.04.756602

**Authors:** Mansi Yadav, Vipul Kumar

## Abstract

Computational safety evaluation of phytochemical libraries commonly relies on models trained on experimentally tested compounds and applied to much larger and chemically diverse collections. An important but often overlooked question is whether predicted compounds are adequately represented within the experimental chemical space underlying these models. Here, we examined this question for the Indian Medicinal Plants, Phytochemistry and Therapeutics (IMPPAT) database using experimentally characterized datasets for three anti-target endpoints: hERG blockade, cytochrome P450 (CYP) inhibition, and pregnane X receptor (PXR) induction. After structure standardization, 14,956 unique IMPPAT compounds were compared with 8,877 hERG, 4,905 CYP, and 4,138 PXR reference compounds using consensus nearest-neighbor similarity based on Morgan and atom-pair fingerprints. At a Tanimoto similarity threshold of 0.80, only 0.92%, 0.30%, and 0.46% of IMPPAT compounds had close measured analogs in the hERG, CYP, and PXR datasets, respectively. Relative to the internal nearest-neighbor distributions of the reference datasets, a typical IMPPAT compound lay near the 1st percentile, indicating substantial chemical-space separation. Compounds with close experimental analogs also showed distinct molecular characteristics, with descriptor-based logistic regression yielding cross-validated AUC values of 0.77–0.80. Plant-level enrichment was markedly reduced after excluding widely distributed metabolites and was further restricted when the nearest measured analog was required to have pIC50 or pEC50 ≥5. The remaining signals were concentrated in a limited number of taxa, including *Rauvolfia*, *Sinopodophyllum/Dysosma*, *Cinchona*, and *Catharanthus roseus*. Overall, most IMPPAT phytochemicals lack close structural representation among currently measured anti-target compounds. Computational safety predictions for these molecules should therefore be interpreted as extrapolations and accompanied by explicit measures of structural support. Absence of a close experimental analog should be regarded as a gap in available evidence rather than evidence of safety.

## 1. Introduction

Drug discovery generally starts with the identification of a biological target whose modulation is expected to provide a therapeutic benefit. However, small molecules often interact with proteins other than their intended target. Such additional interactions can sometimes contribute to useful polypharmacological effects, but they can also lead to unwanted biological responses. These unintended interactions are commonly referred to as off-target activities. Among them, anti-targets represent targets whose modulation is considered undesirable because it can negatively affect efficacy, pharmacokinetics, or safety, and may limit the further development of an otherwise promising compound. Therefore, identifying and minimizing anti-target liabilities at an early stage has become an important part of drug discovery and safety assessment (Recanatini et al. 2004; Thai et al. 2013).

Several anti-targets are routinely considered during lead optimization because of their established associations with toxicity and drug–drug interactions. The human ether-à-go-go-related gene (hERG; KCNH2) potassium channel is a classic example. The unintended hERG blockade can impair cardiac repolarization, prolong the QT interval, and increase the risk of potentially serious ventricular arrhythmias, making hERG liability an important consideration during early drug development (Kalyaanamoorthy and Barakat 2018)(Recanatini et al. 2005). Drug-metabolizing cytochrome P450 (CYP) enzymes are another important source of pharmacokinetic and safety concerns during drug development. Inhibition of major CYP isoforms can reduce the metabolism and clearance of co-administered drugs, leading to increased systemic exposure and a greater risk of adverse drug–drug interactions(Recanatini et al. 2005; Kumar et al. 2012; Nettleton and Einolf 2011). PXR represents a different type of safety concern. As a xenobiotic-sensing nuclear receptor, it controls the expression of several drug-metabolizing enzymes and transporters. Activation of PXR can therefore change the metabolism and disposition of co-administered drugs and may contribute to clinically relevant drug– drug interactions(Sinz 2013; Staudinger et al. 2006). Together, hERG, CYP enzymes, and PXR represent mechanistically different but complementary safety liabilities that are commonly evaluated during preclinical drug development and computational ADMET assessment.

Safety evaluation is especially important for natural products. Plant-derived compounds have long served as valuable starting points for pharmacological research and drug discovery, and medicinal-plant phytochemicals continue to be widely investigated for a range of therapeutic applications. However, products described as natural, herbal, or plant-derived are often assumed to be inherently safe. This assumption can be misleading, because natural origin does not rule out pharmacological or toxicological effects. Phytochemicals may cause direct toxicity, interfere with drug-metabolizing enzymes or transporters, or interact with conventional medicines. In addition, safety information for many herbal products and individual phytochemicals remains much less complete than that available for conventional drugs(Gaston et al. 2020; Shaw et al. 2012; Jordan et al. 2010; Ernst 1998). Therefore, the potential therapeutic activity of a phytochemical should be considered together with its possible safety liabilities rather than assuming safety based only on its natural origin.

Computational ADMET and toxicity prediction provides a practical way to screen large chemical libraries before carrying out experimental testing. Accordingly, QSAR and machine-learning models have been developed for several safety-related endpoints, including hERG blockade, CYP inhibition, PXR activation, hepatotoxicity, and other toxicological properties. However, the reliability of these models depends strongly on the chemical space covered by the experimental data used for model development. In general, a model learns relationships within the molecular and structural space represented in its training set. Predictions for compounds similar to those already present in the training data are therefore closer to interpolation, whereas predictions for structurally different compounds involve increasing levels of extrapolation. Even in such cases, a model may still return a precise numerical prediction, which can give an impression of confidence despite limited experimental support. This issue is addressed through the concept of the applicability domain in QSAR modelling, which defines the region of chemical space where model predictions are considered sufficiently supported by the training data(De et al. 2022; Kar et al. 2018; Jaworska et al. 2005). Recent initiatives such as OpenADMET also aim to improve the accessibility, transparency, and benchmarking of ADMET prediction by providing open experimental datasets and standardized modelling challenges (Fraser et al. 2026).

However, these limitations become particularly important when computational models are applied to phytochemical libraries. Natural products often contain structural features, stereochemical complexity, substitution patterns, and physicochemical properties that differ from those commonly represented in conventional drug-like screening collections. As a result, safety models developed mainly using previously tested drug-like compounds may be applied to plant-derived molecules that are poorly represented in the underlying experimental chemical space. In such cases, the important question is not only what activity value a model predicts, but also whether enough chemically relevant experimental data exist to support that prediction. It is therefore essential to distinguish between predicted safety and lack of experimental coverage. A favourable computational prediction for a phytochemical with little structural similarity to previously tested compounds should not, by itself, be interpreted as evidence of safety(Kar et al. 2018; Jaworska et al. 2005).

The present study examines whether currently available experimental anti-target data adequately cover the chemical space represented by phytochemicals from Indian medicinal plants. Using the IMPPAT database, we address five main questions: (i) what fraction of phytochemicals have close structural analogues among compounds experimentally tested against hERG, CYP enzymes, and PXR; (ii) which regions of phytochemical chemical space are relatively well covered or poorly represented by existing anti-target datasets; (iii) whether phytochemicals with and without close experimental analogues differ in their molecular and structural properties; (iv) whether certain medicinal plants, or their constituent compounds, are disproportionately represented within or outside the experimentally characterized anti-target space; and (v) whether the estimated coverage changes when the analysis is limited to reference compounds showing stronger activity against the corresponding anti-target. By focusing first on chemical-space coverage and applicability domain rather than direct activity prediction, we aim to separate phytochemicals for which existing anti-target data provide meaningful structural support from those for which safety assessment is still limited mainly by insufficient experimental information.

## 2. Materials and Methods

### 2.1 Reference anti-target datasets

The hERG reference set was taken from the ToxTree dataset described by *Arab and Barakat*, containing 8,879 compounds with pIC50 values and the original development/test-set assignment. The dataset was assembled from ChEMBL, PubChem, and literature sources and was originally divided into 8,380 training and 499 test compounds. After structure standardization in the present study, 8,877 unique structures were retained. The dataset does not provide per-measurement uncertainty intervals (Arab et al. 2022).

CYP data were obtained from the *OpenADMET* CYP dataset. The direct-inhibition training table contains 4,905 compounds with pIC50 measurements for CYP1A2, CYP2C9, CYP2D6, and/or CYP3A4, together with 95% confidence intervals and standard deviations derived from the dose-response measurements. Measurements are sparse across isoforms: each compound has at least one measured isoform, whereas only 41 compounds have measurements for all four isoforms (https://huggingface.co/datasets/openadmet/cyp-challenge-train-test).

PXR data were obtained from the *OpenADME*T PXR dataset. The primary assay training table contains 4,139 compounds with pEC50 measurements for PXR induction together with associated standard errors and confidence intervals; 4,138 unique structures remained after structure standardization. Of these, 1,370 compounds also occur in the CYP dataset, and CYP and PXR analyses therefore should not be considered statistically independent (Open ADMET Consortium 2026). The 513-compound test set, released with labels after the challenge (253 compounds in phase 1 and 260 in phase 2), was scored with the same consensus rule and was not added to this reference set.

The three reference datasets were analyzed separately rather than combined because hERG blockade, CYP inhibition, and PXR induction represent mechanistically distinct biological endpoints.

### 2.2 IMPPAT query set and structure processing

IMPPAT contains 189,386 plant-part-compound associations covering 4,010 plants and 17,967 named phytochemicals. Chemical structures were retrieved from PubChem using compound names (Mohanraj et al. 2018). This resulted in 15,119 structure files, from which 14,956 unique standardized structures were retained. During standardization, only the largest covalent fragment was kept, thereby removing salts and other smaller disconnected components, and duplicate structures were identified using the full InChIKey so that stereoisomers remained distinct. Tautomers were not canonicalized, and formal charges were not normalized. A total of 2,835 compound names could not be mapped to chemical structures and were therefore excluded from further analysis. These unmapped compounds were generally sparsely reported: their median occurrence was one plant, and 80% were associated with only a single plant, compared with 58% of the successfully mapped compounds. Thus, the missing structures were concentrated mainly among less frequently reported entries rather than among widely distributed metabolites.

### 2.3 Consensus nearest-neighbor similarity

For each IMPPAT compound, Tanimoto similarity was calculated against all compounds in the three reference datasets using two different 2,048-bit molecular fingerprints: Morgan fingerprints with radius 2 and hashed atom-pair fingerprints (Rogers and Tanimoto 1960)(Landrum, n.d.). For each IMPPAT– reference compound pair, the consensus similarity was calculated by taking the lower of the two fingerprint similarity values. The highest consensus similarity across all compounds in a reference dataset was then assigned as the nearest-neighbor similarity for that IMPPAT compound. This approach ensured that a compound was considered structurally similar only when both fingerprint methods supported the match.

An IMPPAT compound was considered structurally close to an experimentally measured reference compound when its consensus nearest-neighbor Tanimoto similarity was at least 0.80. This threshold was adopted from the supporting analog-pair analysis and was not specifically optimized for the IMPPAT dataset. To examine the effect of the selected cutoff, additional analyses were performed at similarity thresholds of 0.70 and 0.90. We also examined how the results changed when the mean of the two fingerprint similarities was used instead of the minimum.

### 2.4 Uncertainty estimates

The fraction of IMPPAT structures classified as close was summarized with 95% bootstrap confidence intervals using 2,000 resamples of the 14,956 compounds. Mean differences in descriptors between close and far groups were estimated with independent bootstrap resampling of the two groups. For CYP and PXR, the assay confidence interval of the nearest measured neighbor was retained when available. These intervals describe uncertainty in the reference measurement and should not be interpreted as predicted confidence intervals for the corresponding phytochemical.

### 2.5 Descriptor-based separation of close and far compounds

Twenty-four two-dimensional molecular descriptors were calculated to capture size, lipophilicity, polarity, hydrogen-bonding capacity, ring systems, saturation, topological complexity, and the presence of ionizable/basic nitrogen. For each anti-target, L2-regularized logistic regression with balanced class weights was evaluated under five-fold cross-validation. Standardization was performed within each fold. Every descriptor was tested on its own. The eight with the highest cross-validated scores on the full dataset were then combined into pairs and triples, and the full 24-descriptor model was evaluated in the same comparison. Because those eight were chosen using the full dataset, the pair and triple results can be slightly optimistic. The 200 label permutations were applied to the 24- descriptor model and did not repeat that search (Pedregosa et al. 2012; Landrum, n.d.).

### 2.6 Plant-level enrichment analysis

For each medicinal plant, we compared the observed number of structurally close phytochemicals with the number expected under a hypergeometric null distribution. This analysis preserved the total number of compounds reported for each plant while randomly assigning compounds from the IMPPAT collection. The expected distribution was estimated using 20,000 random draws for each distinct number of plant constituents. To account for multiple comparisons across plants, the resulting p-values were adjusted using the Benjamini–Hochberg procedure(Benjamini et al. 2001).

Plant-level enrichment was examined using two approaches. First, the analysis included all structurally close compounds. In the second approach, a compound reported from more than 20 plants was removed from the hit list only. It remained in that plant’s constituent count and in the background pool, where it was treated as a non-hit. Plant list lengths were not reduced. Such compounds may contribute to apparent enrichment across multiple plants without providing information specific to individual plant species. To assess the effect of this filtering criterion, we also repeated the enrichment analysis using alternative limits of 10 and 50 plants.

We further performed a potency-restricted analysis to identify plants containing phytochemicals structurally similar to experimentally active anti-target compounds. For this analysis, only phytochemicals whose nearest measured reference compound had a pIC50 or pEC50 of at least 5, corresponding to an IC50 or EC50 of 10 µM or lower, were retained. This value is a screening threshold used to separate stronger from weaker measured neighbors. Exposure, target context, and assay conditions also determine whether that activity is a clinical liability. For CYP, the highest measured potency among the available isoforms was considered. Plants showing enrichment based on only one retained compound were reported for completeness but were not interpreted as independent biological signals. To examine whether the selection of the nearest neighbor influenced the results, we repeated the analysis by considering the highest measured potency among all reference compounds with a consensus Tanimoto similarity of at least 0.80, rather than restricting the analysis to the single nearest neighbor.

All analyses were performed using Python 3.11, with RDKit 2026.03.6 for molecular structure processing and descriptor calculations and scikit-learn 1.9.1 for machine-learning analyses (Landrum, n.d.)(Pedregosa et al. 2012).

## 3. Results

### 3.1 Reference datasets occupy different internal chemical spaces

The three anti-target reference datasets showed considerable differences in their internal structural similarity and diversity (**Figure 1**). The hERG dataset contained a relatively high proportion of structurally related compounds, with a median nearest-neighbor Tanimoto similarity of 0.79 among 8,877 compounds and 4,427 unique Murcko scaffolds (Landrum, n.d.). In comparison, the CYP and PXR datasets showed lower median nearest-neighbor similarities of 0.45 and 0.39, respectively. The CYP dataset contained 4,519 Murcko scaffolds among 4,905 compounds, while the PXR dataset contained 3,669 scaffolds among 4,138 compounds. These results indicate that the CYP and PXR datasets covered more diverse molecular scaffolds, whereas the hERG dataset contained a larger number of structurally related compounds belonging to similar chemical series. However, compounds across different similarity ranges showed a broad distribution of measured potencies, suggesting that the main difference between these datasets was their structural composition rather than their potency distributions.

**Figure 1.**
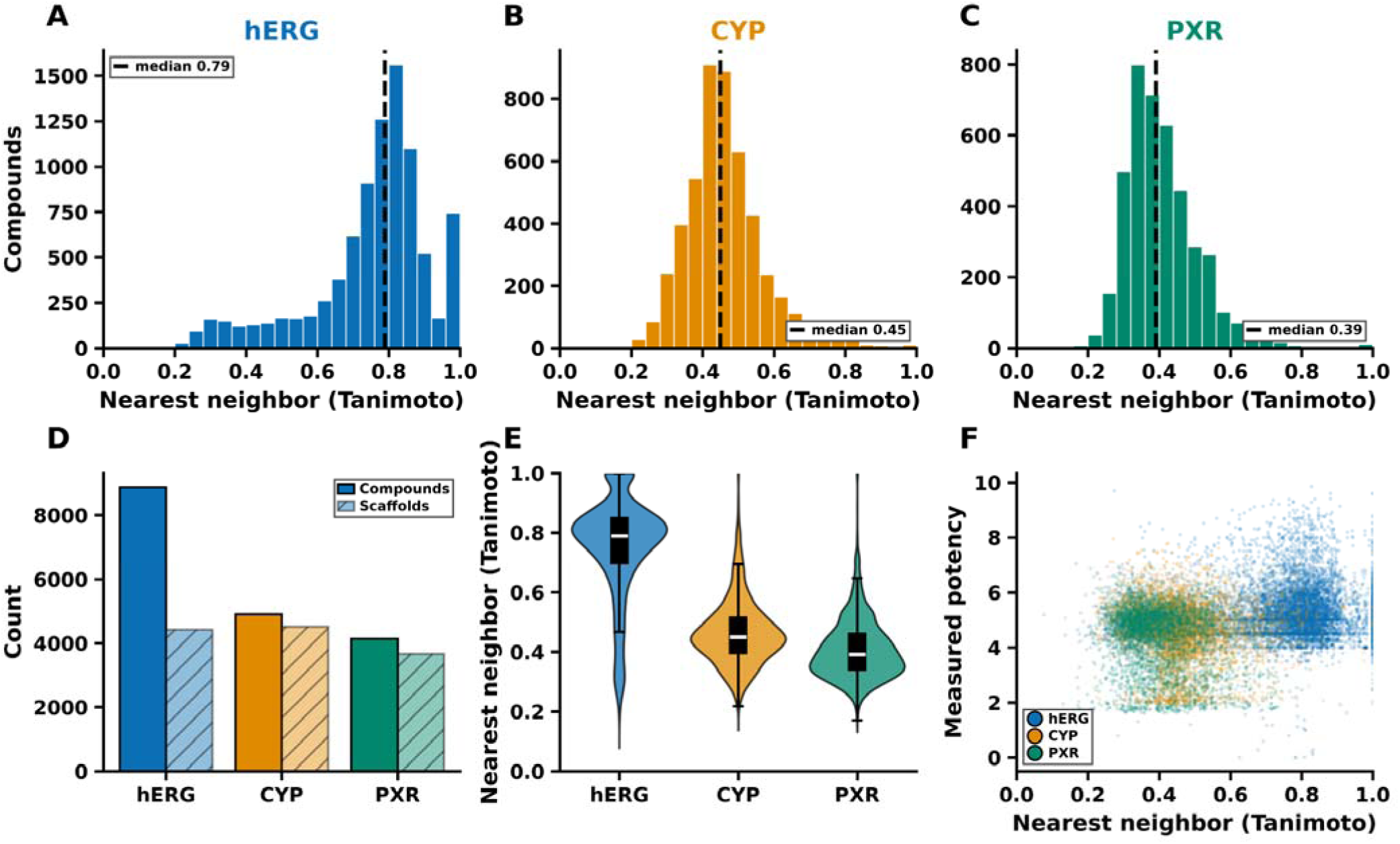
Internal structural diversity of the hERG, CYP, and PXR reference sets. (**A–C**) Distributions of within-set nearest-neighbor Tanimoto similarity, with dashed lines indicating the median. (**D**) Numbers of compounds and *Murcko* scaffolds. (**E**) Violin plots summarizing nearest-neighbor similarity distributions. (**F**) Relationship between nearest-neighbor similarity and measured potency (pIC50 for hERG and CYP; pEC50 for PXR)..

These differences are important when interpreting the structural similarity of IMPPAT phytochemicals to the reference datasets. A low similarity to CYP or PXR reference compounds indicates limited structural resemblance to relatively diverse chemical collections. In contrast, low similarity to the hERG dataset may also reflect the greater representation of particular medicinal-chemistry series among previously tested compounds. Therefore, differences in the internal chemical diversity of the reference datasets should be considered when comparing chemical-space coverage across the three anti-target datasets.

### 3.2 Most IMPPAT phytochemicals are distant from measured anti-target chemistry

The majority of IMPPAT phytochemicals showed low structural similarity to compounds in the three experimentally characterized anti-target datasets (**Figure 2**). The median nearest-neighbor Tanimoto similarity was approximately 0.28 for hERG and 0.26 for CYP, while PXR showed a similarly low similarity distribution. For all three datasets, the cumulative similarity distributions approached 1.0 well before reaching the threshold of 0.80, indicating that only a small fraction of IMPPAT compounds had highly similar experimental reference compounds. In addition, comparison with the 513 unblinded PXR test compounds showed a maximum consensus similarity of only 0.42, indicating limited structural overlap with this additional set (**Figure 2E**).These 513 compounds are the post-challenge unblinded test set and were not part of the 4,138-compound reference collection.

**Figure 2.**
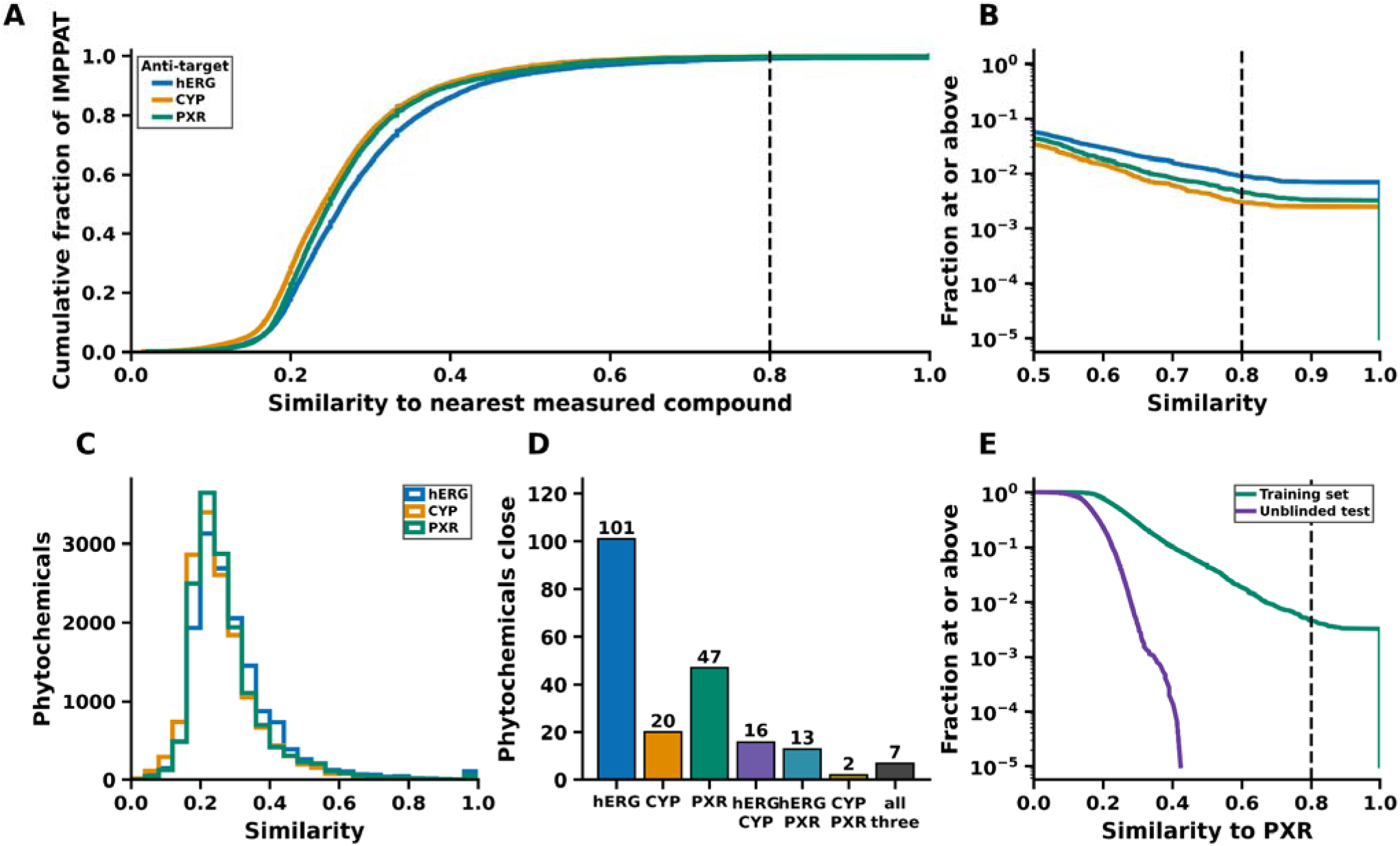
Similarity of IMPPAT phytochemicals to measured hERG, CYP, and PXR compounds. (**A**) Cumulative fraction of the 14,956 IMPPAT structures against similarity to the nearest measured compound. (**B**) The same comparison on the upper tail, as the fraction of IMPPAT at or above a given similarity. (**C**) Histogram of that similarity. (**D**) Phytochemicals with consensus similarity of at least 0.80, split into mutually exclusive groups: hERG only, CYP only, PXR only, hERG and CYP, hERG and PXR, CYP and PXR, and all three. (**E**) Fraction of IMPPAT at or above a given similarity to PXR, for the training set and the 513 unblinded test compounds.

At a consensus Tanimoto similarity threshold of ≥0.80, only 137 of the 14,956 IMPPAT phytochemicals had close structural analogues in the hERG dataset, representing 0.92% of the library (95% CI: 0.78–1.07%). Similarly, only 45 compounds (0.30%, 95% CI: 0.22–0.39%) were close to the CYP reference dataset, while 69 compounds (0.46%, 95% CI: 0.35–0.58%) showed close similarity to the PXR dataset. The overlap between endpoints was even smaller. Only 23 phytochemicals had close analogues in both the hERG and CYP datasets, 20 in both hERG and PXR, and 9 in both CYP and PXR. Notably, only 7 compounds showed close structural similarity to reference compounds across all three anti-target datasets (**Table 1**). All nine IMPPAT phytochemicals showing close structural similarity to both the CYP and PXR datasets had their nearest neighbors among the 1,370 compounds shared between the two reference datasets. Thus, the observed CYP–PXR overlap mainly reflects the presence of common reference compounds rather than independent structural matches across the two datasets.

**Table1.** Calculated Tanimoto similarity between IMPPAT database compounds with reference sets.

| Anti-target | Reference compounds | Internal median NN | Close at 0.80 | Fraction (95% CI) |
| --- | --- | --- | --- | --- |
| hERG | 8,877 | 0.79 | 137 | 0.92% (0.78–1.07) |
| CYP | 4,905 | 0.45 | 45 | 0.30% (0.22–0.39) |
| PXR | 4,138 | 0.39 | 69 | 0.46% (0.35–0.58) |
| hERG ∩ CYP | — | — | 23 | 0.15% |
| hERG ∩ PXR | — | — | 20 | 0.13% |
| CYP ∩ PXR | — | — | 9 | 0.06% |
| All three | — | — | 7 | 0.05% |

### 3.3 Requiring fingerprint agreement removes high-similarity artifacts

At all tested similarity thresholds, the two-fingerprint consensus approach identified fewer structurally close IMPPAT phytochemicals than Morgan fingerprints alone (**Figure 3**). This difference became more pronounced at higher similarity thresholds. For example, at a Tanimoto cutoff of 0.90, Morgan fingerprints identified 140 phytochemicals with close structural similarity to the hERG reference compounds, whereas the consensus approach reduced this number to 106. Similar trends were observed for CYP and PXR. At the 0.80 threshold, the number of close compounds decreased from 212 to 137 for hERG, 85 to 45 for CYP, and 147 to 69 for PXR after applying the consensus criterion. We also compared this fixed threshold with the median nearest-neighbor similarity within each reference dataset to account for differences in their internal structural diversity (**Figure 3D**).

**Figure 3.**
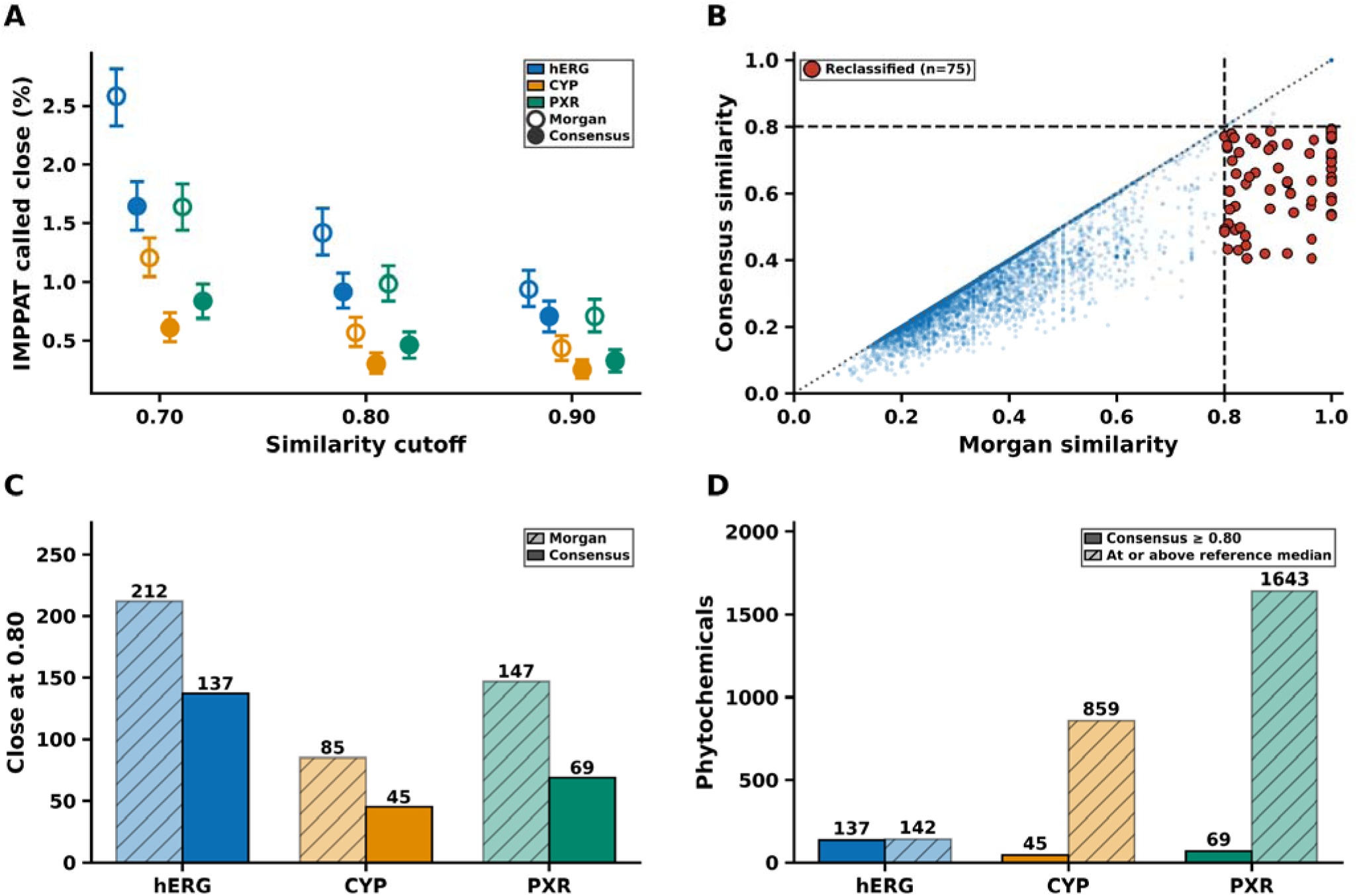
Robustness of the close-compound definition to similarity cutoff and fingerprint representation. (**A**) Percentage of IMPPAT compounds called close at Tanimoto cutoffs of 0.70, 0.80, and 0.90 using Morgan fingerprints or the Morgan/atom-pair consensus. (**B**) Relationship between Morgan and consensus similarity; red points indicate compounds reclassified by the consensus rule. (**C**) Counts of compounds called close at the 0.80 threshold under the two definitions. (D) Numbers of phytochemicals with consensus similarity of at least 0.80, compared with the numbers whose consensus similarity reaches that reference set’s own median nearest-neighbor similarity.

The compounds excluded by the consensus approach mainly included structures with similar local molecular environments but differences in their overall topology, particularly long aliphatic chains. These findings indicate that the estimated chemical-space overlap between IMPPAT phytochemicals and experimentally characterized anti-target compounds depends on the fingerprint representation used. Therefore, combining Morgan fingerprints with a topologically sensitive atom-pair fingerprint provides a more conservative assessment of structural similarity and helps reduce potentially misleading high-similarity matches.

### 3.4 Close phytochemicals form a chemically distinguishable minority

Although only a small fraction of IMPPAT phytochemicals showed close structural similarity to the reference datasets, these compounds differed in their molecular properties from the remaining phytochemicals (**Figure 4**). Logistic regression models using 24 two-dimensional molecular descriptors distinguished structurally close and distant compounds, with cross-validated AUC values of 0.77 for hERG, 0.73 for CYP, and 0.80 for PXR. In comparison, the 200 label-permutation controls produced AUC values close to 0.50 (0.49–0.50), and none achieved the performance of the corresponding models trained using the original labels.

**Figure 4.**
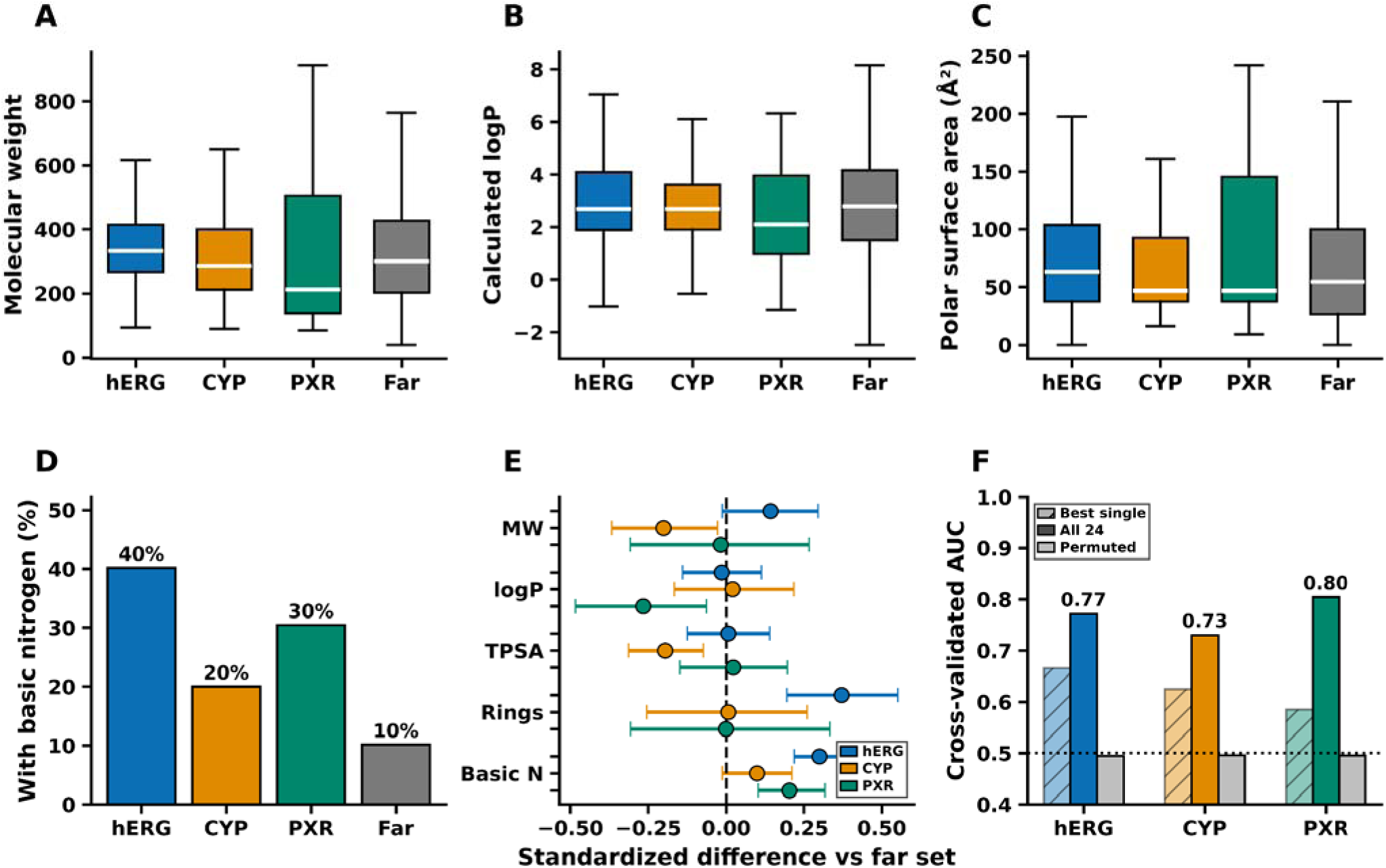
Molecular-property differences between phytochemicals close to and far from the measured anti-target sets. (**A–C**) Distributions of molecular weight, calculated logP, and polar surface area. (**D**) Fraction containing basic nitrogen. (**E**) Descriptor differences divided by each descriptor’s standard deviation, with uncertainty intervals. Basic nitrogen is shown as a difference in proportion. (**F**) Five-fold cross-validated AUC for the best single descriptor, the full 24-descriptor model, and label-permuted controls.

Individual molecular descriptors or combinations of a few descriptors did not perform as well as the complete 24-descriptor model. Among individual descriptors, aromatic ring count showed the highest AUC for hERG (0.67), fraction of sp³ carbons for CYP (0.62), and Bertz complexity for PXR (0.59). The best models using three descriptors achieved AUC values of 0.75, 0.65, and 0.70 for hERG, CYP, and PXR, respectively. These results suggest that the structural differences between close and distant phytochemicals are associated with a combination of molecular properties rather than any single descriptor.

Further examination of molecular descriptors revealed differences between the two groups. Phytochemicals close to the hERG reference dataset had a higher average ring count than distant compounds (mean difference 0.86, 95% CI: 0.45–1.28) and a greater proportion of compounds containing basic or ionizable nitrogen (difference 0.30, 95% CI: 0.22–0.38). These characteristics are consistent with structural features commonly observed among hERG-binding compounds, particularly aromatic ring systems and protonatable amines. For PXR, close phytochemicals showed lower lipophilicity, with a mean logP difference of −0.77 (95% CI: −1.40 to −0.18), and a higher prevalence of basic nitrogen (difference 0.20, 95% CI: 0.10–0.32). Similarly, CYP-close phytochemicals generally had lower molecular weights (mean difference −42 Da, 95% CI: −77 to −6) and lower topological polar surface areas (mean difference −17 Å², 95% CI: −27 to −6) compared with distant compounds.

However, the observed AUC values of 0.73–0.80 indicate that considerable overlap remains between the molecular properties of close and distant phytochemicals. Therefore, these differences should be interpreted as general trends across the compound groups rather than as definitive characteristics of individual molecules. Importantly, because the close and distant groups were defined based on their structural similarity to experimentally measured reference compounds, the descriptor analysis primarily describes the types of chemical structures represented in the existing anti-target datasets. It should not be considered independent evidence that the close phytochemicals constitute a distinct biological or pharmacological class.

### 3.5 Plant-level enrichment is largely driven by ubiquitous compounds unless prevalence is controlled

When all structurally close phytochemicals were included in the enrichment analysis, a large number of medicinal plants showed statistically significant enrichment (q < 0.05), with 566 plants for hERG, 864 for CYP, and 860 for PXR. However, these numbers decreased substantially after excluding compounds reported in more than 20 plants. Following this restriction, only 12 plants remained significantly enriched for hERG, 6 for CYP, and 4 for PXR (**Figure 5**). To examine whether these results depended on the selected prevalence cutoff, we repeated the analysis using limits of 10 and 50 plants. At a cutoff of 10 plants, the numbers of significantly enriched species were 8, 3, and 7 for hERG, CYP, and PXR, respectively, while a cutoff of 50 plants resulted in 16, 6, and 5 significant species. These results indicate that the large reduction in plant-level enrichment was consistent across different prevalence thresholds and was not specific to the cutoff of 20 plants.

**Figure 5.**
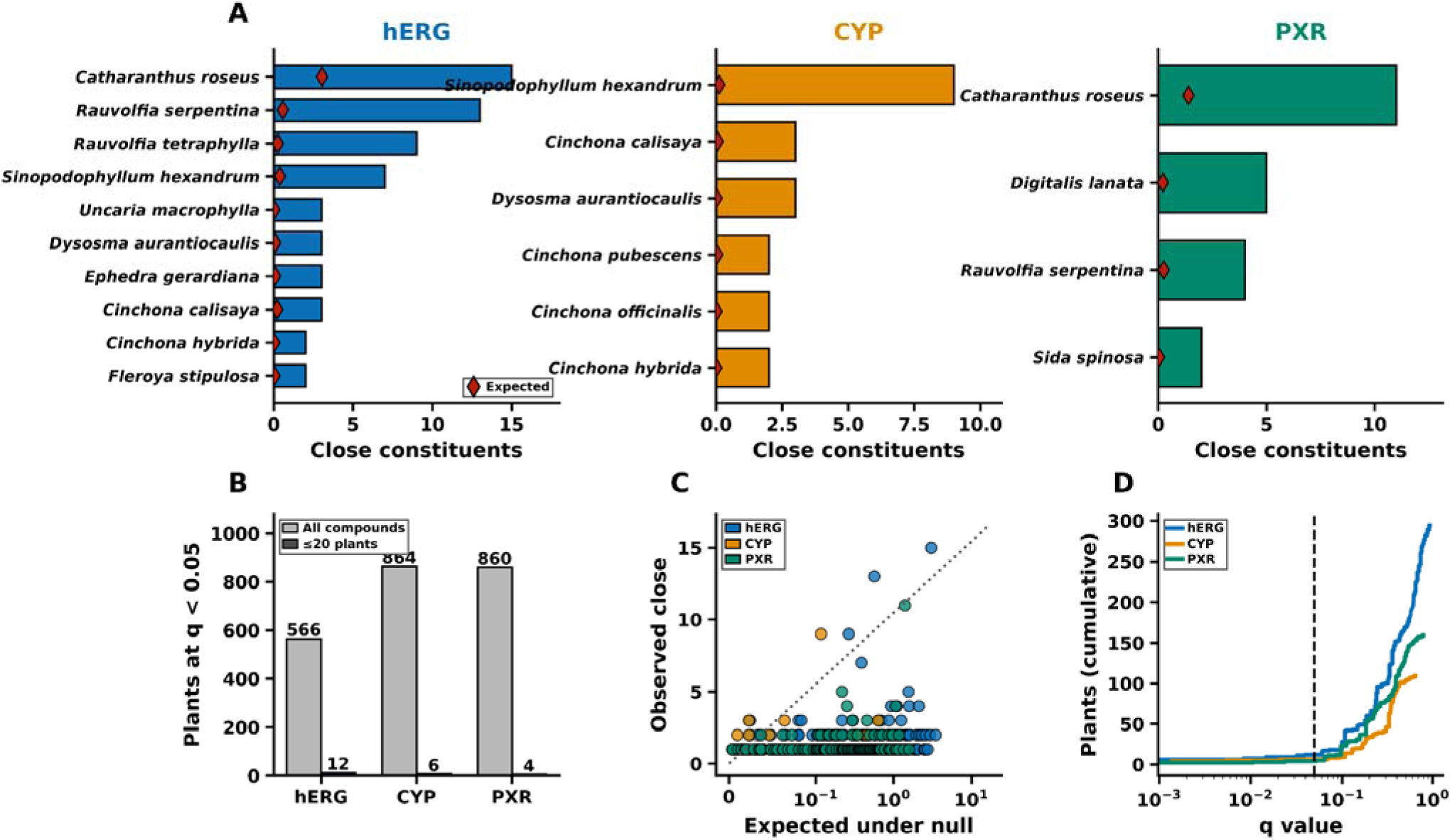
Plant-level enrichment of phytochemicals structurally close to measured anti-target compounds. (**A**) Plants with the largest numbers of close constituents after excluding compounds recorded from more than 20 plants; red diamonds show counts expected under the null. (**B**) Number of significant plants before and after prevalence restriction. (**C**) Observed versus null-expected numbers of close constituents. (**D**) Cumulative number of plants as a function of q value.

Among the plants showing significant enrichment for hERG, the prominent species included *Catharanthus roseus*, *Rauvolfia serpentina, Rauvolfia tetraphylla, Sinopodophyllum hexandrum, Dysosma aurantiocaulis, Uncaria macrophylla*, and *Ephedra gerardiana*. For CYP, enrichment was mainly observed in *Sinopodophyllum, Dysosma,* and several *Cinchona* species. In comparison, PXR enrichment was associated with *Digitalis lanata, Catharanthus roseus,* and *Rauvolfia serpentina*. These plants are known to contain different classes of pharmacologically active compounds, including monoterpene indole alkaloids, aryltetralin lignans, quinoline alkaloids, cardiac glycosides, phenylethylamines (**Table 2**).

**Table2.** Plants enriched for phytochemicals close to a measured anti-target compound.

| Anti-target | Plant | Constituents | Close | Expected | q |
| --- | --- | --- | --- | --- | --- |
| hERG | *Catharanthus roseus* | 384 | 15 | 3.05 | <0.001 |
| hERG | *Rauvolfia serpentina* | 70 | 13 | 0.57 | <0.001 |
| hERG | *Rauvolfia tetraphylla* | 33 | 9 | 0.27 | <0.001 |
| hERG | *Sinopodophyllum hexandrum* | 48 | 7 | 0.39 | <0.001 |
| hERG | *Dysosma aurantiocaulis* | 10 | 3 | 0.08 | <0.001 |
| hERG | *Uncaria macrophylla* | 3 | 3 | 0.02 | <0.001 |
| hERG | *Ephedra* | 11 | 3 | 0.08 | 0.008 |
|  | *gerardiana* |  |  |  |  |
| CYP | *Sinopodophyllum hexandrum* | 48 | 9 | 0.12 | <0.001 |
| CYP | *Dysosma aurantiocaulis* | 10 | 3 | 0.02 | <0.001 |
| CYP | *Cinchona calisaya* | 27 | 3 | 0.06 | 0.001 |
| CYP | *Cinchona hybrida* | 4 | 2 | 0.01 | 0.001 |
| PXR | *Digitalis lanata* | 60 | 5 | 0.22 | <0.001 |
| PXR | *Catharanthus roseus* | 384 | 11 | 1.40 | <0.001 |
| PXR | *Rauvolfia serpentina* | 70 | 4 | 0.25 | 0.011 |

The marked difference between the unrestricted and prevalence-restricted analyses suggests that much of the initial plant-level enrichment was driven by widely distributed metabolites rather than compounds specific to particular plants. Excluding these common metabolites therefore helped identify plant species whose reported constituents showed greater structural similarity to experimentally characterized anti-target compounds. Interestingly, the analysis identified plants known to contain specialized classes of bioactive compounds, even though no experimental activity information for the IMPPAT phytochemicals was used. This provides an internal check that the similarity-based approach can identify recognizable chemical groups when they are represented in the reference datasets. In contrast, the absence of enrichment for other plant species should not be interpreted as evidence of safety, as their constituent compounds may simply lack close structural representation in the available anti-target datasets.

### 3.6 Potency filtering further narrows the biologically interpretable set

Structural similarity to an experimentally tested compound does not necessarily indicate that the reference compound has strong activity against the corresponding anti-target. Among the 137 hERG-close, 45 CYP-close, and 69 PXR-close phytochemicals, only 51, 16, and 11 compounds, respectively, had a nearest measured neighbor with a pIC50 or pEC50 of at least 5. The median potency of the nearest measured neighbors was 4.85 for hERG, 4.70 for CYP, and 2.50 for PXR, suggesting that most PXR-close phytochemicals were structurally similar to relatively weak inducers. We also examined whether these results changed when the most potent reference compound with a consensus Tanimoto similarity of at least 0.80 was considered instead of the single nearest neighbor. The number of compounds remained unchanged for hERG (51) and PXR (11), while the CYP count increased from 16 to 19. The three additional compounds were quinine, quinidine, and epiquinine, whose nearest CYP neighbors had pIC50 values below 5, whereas another sufficiently similar reference compound had a pIC50 of 7.47. However, under the original nearest-neighbor criterion, Cinchona species were no longer significantly enriched for CYP after potency filtering (**Table 3**).

**Table 3:** Plants that remain enriched after the nearest measured neighbor is required to be potent.

| Anti-target | Close | Neighbor $\geq 5$ | Plants at $q < 0.05$ with $\geq 2$ retained constituents |
| --- | --- | --- | --- |
| hERG | 137 | 51 | *Rauvolfia serpentina*; *R. tetraphylla*; *Sinopodophyllum hexandrum*; *Dysosma aurantiocaulis*; *Cinchona calisaya*; *Alangium salviifolium*; *Carapichea ipecacuanha*; *Croton tiglium*; *Camellia sinensis* |
| CYP | 45 | 16 | *Sinopodophyllum hexandrum*; *Dysosma aurantiocaulis* |
| PXR | 69 | 11 | *Catharanthus roseus* |

Applying the potency requirement also changed the plants identified in the enrichment analysis. For hERG, significant enrichment involving multiple constituents was retained in *Rauvolfia serpentina*, *R. tetraphylla, Sinopodophyllum hexandrum, Dysosma aurantiocaulis*, and *Cinchona calisaya,* along with several additional species. For CYP, only *Sinopodophyllum hexandrum and Dysosma aurantiocaulis* retained significant enrichment involving multiple constituents. In contrast, *Catharanthus roseus* was the main enriched species for PXR, with nine of the eleven phytochemicals having potent nearest neighbors associated with vinca alkaloid chemistry and matching an experimentally measured reference compound with a pEC50 of 5.64.

The observed numbers of potent-neighbor phytochemicals were considerably higher than expected for several enriched plants. For hERG, *Rauvolfia serpentina* contained five such compounds compared with an expected 0.22, while *R. tetraphylla* contained four compared with 0.10. Similarly, *Sinopodophyllum hexandrum* contained seven compared with an expected 0.15, *Dysosma aurantiocaulis* contained three compared with 0.03, *Cinchona calisaya* contained three compared with 0.09, and A*langium salviifolium* contained three compared with 0.28. Additional enriched plants included *Carapichea ipecacuanha, Croton tiglium*, and *Camellia sinensis*. For CYP, *Sinopodophyllum hexandrum* had seven potent-neighbor compounds compared with an expected 0.04, while *Dysosma aurantiocaulis* had three compared with 0.01. For PXR, *Catharanthus roseus* contained nine such compounds compared with an expected 0.28.

A small number of additional plants, particularly several Sophora species, showed significant enrichment based on a single compound, cytisine, whose nearest measured PXR neighbor had a pEC50 of 6.09. However, a single compound can produce a statistically significant enrichment value when the total number of reported constituents for a plant is small. Therefore, these single-compound signals were not considered independent evidence of plant-level enrichment. Overall, the potency-restricted analysis identified a smaller group of medicinal plants whose constituents showed structural similarity to experimentally active anti-target compounds, while also demonstrating that structural similarity alone does not necessarily indicate a strong pharmacological liability.

### 3.7 A small number of close matches are chemically plausible but biologically questionable records

All 206 unique IMPPAT structures showing a consensus Tanimoto similarity of at least 0.80 to one or more reference datasets were examined and classified into four categories: putative phytochemical constituents (170), xenobiotics or drugs reported as plant constituents (20), industrial or combustion-related contaminants (8), and compounds with uncertain origins (8). Some potentially questionable records included quinacrine reported in Brucea javanica, dipyridamole in Apium graveolens, cyproheptadine hydrochloride in Melia azedarach, triphenylphosphine in Mukia maderaspatana, and fluoranthene in Cichorium intybus. These labels are a plausibility assessment of the recorded plant occurrence. They are not experimental verification that a compound is a genuine plant constituent These compounds may reflect database annotation errors, contamination, or limitations in the original source records rather than genuine plant-derived constituents.

To examine the influence of these records on the overall results, we repeated the coverage analysis after excluding the 28 compounds classified as xenobiotics or contaminants. This reduced the number of structurally close phytochemicals from 137 to 113 for hERG, 45 to 40 for CYP, and 69 to 65 for PXR. The eight compounds with uncertain origins, including several mammalian steroids, were retained because their status could not be established confidently. Importantly, the main coverage results reported in the preceding sections include all close structures, including potentially questionable records. Thus, excluding these compounds further reduces the estimated structural coverage rather than increasing it, supporting the observation that only a small fraction of IMPPAT structures have close experimental analogues in the available anti-target datasets.

### 3.8 The coverage does not depend on one absolute cutoff

To examine whether the estimated chemical-space coverage depended on the selected similarity threshold, we compared two approaches: a fixed consensus Tanimoto similarity cutoff of 0.80 and a dataset-specific cutoff based on the median nearest-neighbor similarity within each reference set (**Figure 3D**). At the fixed threshold of 0.80, only 137, 45, and 69 IMPPAT phytochemicals were classified as structurally close to the hERG, CYP, and PXR datasets, respectively. When the median nearest-neighbor similarity of each reference dataset was used instead, these numbers increased to 142 for hERG, 859 for CYP, and 1,643 for PXR. The difference was small for hERG because its internal median similarity was approximately 0.80. In contrast, the CYP and PXR datasets had considerably lower internal median similarities, indicating that the fixed threshold of 0.80 was more stringent for these structurally diverse datasets. Thus, the conclusion that more than 99% of IMPPAT compounds lack close structural analogues depends partly on the selected absolute similarity cutoff.

Despite these differences in coverage estimates, most IMPPAT phytochemicals remained structurally distant when compared with the internal similarity distributions of the reference datasets. The median consensus similarity of IMPPAT compounds corresponded to approximately the 1st percentile of the within-reference nearest-neighbor distributions for all three endpoints. Furthermore, 80%, 76%, and 65% of IMPPAT compounds had similarity values below the 5th percentile of the corresponding hERG, CYP, and PXR reference distributions, respectively. These findings indicate that the low structural representation of typical IMPPAT compounds is not solely a consequence of using the fixed 0.80 threshold, although the estimated fraction of structurally close compounds changes with the definition of coverage. Each reference-set median describes the internal diversity of that dataset. It has not been shown to be a boundary for reliable prediction.

We also examined CYP chemical-space coverage separately for each isoform, considering only reference compounds with experimental measurements for the corresponding enzyme. At a consensus Tanimoto threshold of 0.80, 7 IMPPAT phytochemicals had close analogues among 1,412 CYP1A2 reference compounds, 25 among 1,285 CYP2C9 compounds, 11 among 1,493 CYP2D6 compounds, and 21 among 2,335 CYP3A4 compounds. The combined set of unique phytochemicals identified across these four isoforms contained 45 structures, matching the number obtained from the pooled CYP analysis. This confirms that the pooled analysis captures the union of isoform-specific structural matches, while the individual isoform results provide a more detailed view of experimental chemical-space coverage.

## 4. Discussion

The present study shows that more than 99% of the IMPPAT phytochemicals examined here do not have a close structural analogue in any individual hERG, CYP, or PXR reference dataset at a consensus Tanimoto similarity threshold of 0.80. This finding has important implications for computational safety assessment of natural products. Although machine-learning models can generate activity predictions for a large number of compounds, the experimental data used to develop these models may not adequately represent the chemical structures being evaluated. Consequently, predictions for such compounds may have limited experimental support and should not be interpreted with the same confidence as predictions for compounds represented in the training data. This observation was also supported by comparisons with the internal similarity distributions of the reference datasets. The median similarity of IMPPAT compounds was approximately at the 1st percentile of the corresponding within-reference nearest-neighbor distributions. Furthermore, 80%, 76%, and 65% of IMPPAT phytochemicals fell below the 5th percentile of the hERG, CYP, and PXR reference distributions, respectively (**Figure 3D**).

Additional analyses using the hERG reference dataset further demonstrated the importance of distinguishing structural similarity from prediction reliability (**Figure S1**). In the cluster-holdout evaluation, prediction errors generally decreased as the similarity to the nearest training compound approached a Tanimoto value of 0.80. However, this relationship was not observed in the published external test set. Even among the 29 test compounds having a training-set neighbor with similarity of at least 0.80, the median absolute prediction error remained 0.50 log units. These findings indicate that structural similarity alone does not guarantee accurate predictions.

These observations do not suggest that computational prediction models are unsuitable for natural-product safety assessment. Rather, they emphasize the importance of considering the applicability domain when interpreting their predictions. The comparisons in this study are to the three experimental collections assembled here, and the hERG collection includes both the published development compounds and the external test compounds. Coverage in a different computational model still has to be judged against that model’s own training set. For phytochemicals with close experimental analogues, the identity and measured activity of the reference compounds can provide useful information for interpreting potential anti-target liabilities. However, even among the structurally close compounds, only 51 hERG, 16 CYP, and 11 PXR phytochemicals had a nearest measured neighbor with a pIC50 or pEC50 of at least 5. Therefore, the presence of a close structural analogue and the strength of its measured activity should be considered separately, because structural similarity does not necessarily indicate a strong pharmacological liability.

The internal chemical diversity of the reference datasets is another important factor affecting the interpretation of chemical-space coverage. The hERG dataset contained a relatively large number of structurally related compounds, reflecting the concentration of experimental measurements within particular medicinal-chemistry series. In contrast, the CYP and PXR datasets covered a broader range of molecular scaffolds. As a result, low structural similarity to CYP or PXR reference compounds indicates separation from relatively diverse experimental collections, whereas low similarity to the hERG dataset may partly reflect the greater representation of specific chemical series. These differences also explain why the same absolute Tanimoto threshold does not provide an equivalent measure of chemical-space coverage across all three endpoints. Therefore, both absolute similarity and the internal diversity of the reference datasets should be considered when evaluating structural representation.

The comparison between Morgan and atom-pair fingerprints further showed that chemical-space coverage depends on the molecular representation used. Morgan fingerprints are commonly used in cheminformatics and machine-learning applications because they efficiently capture local molecular environments. However, high similarity based on Morgan fingerprints alone may not always reflect close structural resemblance, particularly when compounds differ in their overall topology. In the present analysis, combining Morgan and atom-pair fingerprints reduced several potentially misleading matches involving long aliphatic chains and other topologically different structures. This difference became more pronounced at higher similarity thresholds, where structural matches are more likely to be interpreted as close analogues. Thus, using complementary molecular fingerprints can provide a more conservative assessment of chemical similarity and reduce dependence on a single representation.

The descriptor analysis also revealed differences between IMPPAT phytochemicals with and without close experimental analogues. Compounds close to the hERG reference dataset generally contained more rings and were more likely to possess basic or ionizable nitrogen, consistent with structural features commonly associated with hERG-binding compounds. CYP-close phytochemicals tended to have lower molecular weights and polar surface areas, whereas PXR-close compounds showed differences in lipophilicity, nitrogen content, and molecular complexity. However, the descriptor-based models achieved AUC values below 0.81, indicating considerable overlap between the two groups. Furthermore, because structural similarity to the reference datasets was used to define the close and distant groups, these results primarily describe the types of chemical structures represented in existing experimental collections rather than identifying distinct biological classes. The observed differences therefore suggest that chemical-space representation depends on multiple molecular properties and cannot be adequately described using a single physicochemical descriptor.

At the medicinal-plant level, the presence of widely distributed metabolites had a substantial influence on the enrichment results. When all structurally close compounds were included, hundreds of plants appeared significantly enriched. However, after excluding compounds reported in more than 20 plants, the number of enriched species decreased considerably. The remaining plants included *Rauvolfia, Catharanthus, Sinopodophyllum, Dysosma, and Cinchona,* which are known to contain specialized alkaloid or lignan scaffolds. The identification of these plants based only on structural similarity, without using their experimental activity information, provides an internal check that the approach can identify recognizable chemical groups represented in the reference datasets. Nevertheless, enrichment should not be interpreted as direct evidence of toxicity associated with these plants. IMPPAT records the reported occurrence of phytochemicals but does not provide sufficient information about their abundance, exposure, bioavailability, metabolism, or formulation, all of which can influence biological and toxicological effects.

The potency-restricted analysis further demonstrated the importance of considering structural similarity and measured activity separately. Several medicinal plants that were enriched based on structural similarity alone were no longer significantly enriched when the nearest measured analog was required to have a pIC50 or pEC50 of at least 5. This effect was particularly evident for PXR, where the remaining enrichment was mainly associated with *Catharanthus roseus*. These findings suggest that chemical-space coverage and potential pharmacological liability can be evaluated in two stages. First, it is necessary to determine whether a phytochemical has close experimental analogues in the available reference datasets. Second, the measured activities of these analogues can be examined to understand whether they provide evidence relevant to potential anti-target activity. However, the activity of a structurally related compound should not be treated as a direct measurement of the phytochemical itself.

Several limitations of this study should also be considered. First, chemical structures were retrieved from PubChem using compound names, and 2,835 IMPPAT entries could not be mapped successfully. Therefore, the analysis does not include the complete IMPPAT collection and may be affected by structure-retrieval bias. The unmapped names were predominantly associated with fewer plants, with 80% reported from only one plant compared with 58% of mapped names. This suggests that missing structures were concentrated among less frequently reported compounds, although their exclusion may still influence the estimated coverage. Second, IMPPAT is based on phytochemical information reported in the literature, and extensively studied plants may have longer lists of identified constituents than less investigated species. Although the enrichment analysis preserved the number of reported constituents for each plant and excluded widely distributed metabolites, these procedures cannot completely eliminate reporting and ascertainment biases.

Another limitation is the use of structural similarity thresholds to define chemical-space coverage. The Tanimoto cutoff of 0.80 was used as an operational criterion rather than a strict biological boundary, and changing the threshold naturally affected the number of compounds classified as structurally close. To address this limitation, we examined alternative thresholds and compared IMPPAT similarity values with the internal nearest-neighbor distributions of the reference datasets. Although the absolute coverage estimates varied, the typical IMPPAT compound remained within the low-similarity tail of these distributions (**Figure 3D**). Furthermore, the two-dimensional fingerprints used here cannot fully represent stereochemical differences, conformational preferences, protonation states, or three-dimensional interactions involved in molecular recognition. The reference datasets also differ in their experimental designs, assay conditions, and uncertainty estimates, while the CYP and PXR datasets share a substantial number of compounds. These factors should be considered when interpreting differences between endpoints.

Importantly, the absence of a close measured analogue does not imply that a phytochemical is biologically inactive or safe. A compound that is structurally distant from available reference molecules may still inhibit hERG or CYP enzymes or activate PXR. The present analysis identifies limitations in the structural representation of phytochemicals within existing experimental datasets, rather than establishing their biological activities or safety profiles.

Overall, these findings highlight the need to consider experimental chemical-space support when applying computational safety models to natural-product libraries. Activity predictions would be more informative if accompanied by a measure indicating how closely a query compound resembles molecules represented in the corresponding experimental dataset. When close measured analogues are available, their identities, activities, and associated experimental uncertainties can provide additional context for interpreting predictions. In contrast, predictions for structurally distant compounds should be recognized as extrapolations with limited direct experimental precedent. Incorporating such information into computational screening would help distinguish predictions supported by existing measurements from those requiring further experimental investigation.

## 5. Conclusion

Across 14,956 IMPPAT phytochemical structures compared with three experimentally characterized anti-target datasets, fewer than 1% of compounds had close structural analogues in any individual hERG, CYP, or PXR reference dataset at a consensus Tanimoto similarity threshold of 0.80. Even when similarity was evaluated relative to the internal chemical diversity of each reference dataset, a typical IMPPAT compound remained near the 1st percentile of the corresponding nearest-neighbor similarity distribution, indicating limited structural representation. After accounting for widely distributed metabolites and considering the measured potency of the nearest experimental analogues, the structurally represented compounds were mainly associated with a small number of plant taxa, including Rauvolfia, Sinopodophyllum, Dysosma, and Catharanthus roseus. These findings suggest that the available hERG, CYP, and PXR experimental datasets provide close structural precedent for only a small fraction of the phytochemicals represented in IMPPAT. For most compounds, the major limitation is the lack of closely related experimental measurements rather than demonstrated absence of anti-target activity. These results describe representation in the reference collections used here. Applicability of a specific computational model still has to be assessed against that model’s own training set. Therefore, computational safety predictions for such phytochemicals should be interpreted in the context of their structural similarity to experimentally tested compounds. Limited experimental coverage should be clearly reported as a data gap rather than being overlooked when interpreting numerical predictions or considered evidence of safety.

## Author Contribution

Conceptualization: V.K; Methodology: M.Y, V.K.; Software: M.Y., V.K..; Formal Analysis, M.Y., V.K; Original Draft Preparation: V.K.; Writing, Review and Editing: M.Y., V.K; Supervision: V.K. Authors have read and agreed to the published version of the manuscript.

## Acknowledgments

V.K and M.Y acknowledge the Ramanujan Universe High Performance Computing Facility at Jaypee Institute of Information Technology for providing computational resources used in this research.

## Supplementary information

**Figure S1**. Prediction error against distance to the training set (hERG). Under cluster holdout, Gaussian-process RMSE falls from 0.86 for compounds whose nearest training neighbor is below Tanimoto 0.30 to 0.60 in the 0.70-0.80 bin. On the published external test split the relationship does not hold. Binned RMSE is 0.91, 1.07, 0.79, 0.76, 1.09, 0.89, and 0.68 across ascending similarity bins, and the 29 test compounds with a training neighbor at 0.80 or above retain a median absolute error of 0.50 log units.

