## Supplementary figures and images for "Anti-target chemical-space coverage of Indian medicinal plant phytochemicals"

### Supplementary Figure S1

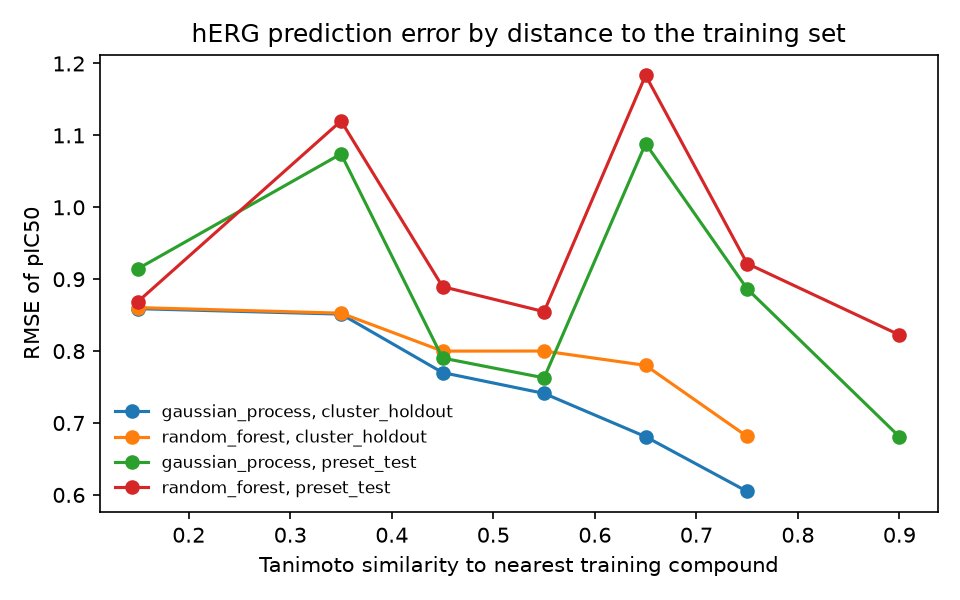
